# Network approach reveals preferential T-cell and macrophage association with α-linked β-cells in early stage of insulitis in NOD mice

**DOI:** 10.1101/2024.05.06.592831

**Authors:** Nirmala V. Balasenthilkumaran, Jennifer C. Whitesell, Laura Pyle, Rachel Friedman, Vira Kravets

## Abstract

One of the challenges in studying islet inflammation – insulitis – is that it is a transient phenomenon. Traditional reporting of the insulitis progression is based on cumulative, donor-averaged values of leucocyte density in the vicinity of pancreatic islets, that hinders intra- and inter-islet heterogeneity of disease progression. Here, we aimed to understand why insulitis is non-uniform, often with peri-insulitis lesions formed on one side of an islet. To achieve this, we demonstrated applicability of network theory in detangling intra-islet multi-cellular interactions during insulitis. Specifically, we asked the question “what is unique about regions of the islet which interact with immune cells first”. This study utilized the non-obese diabetic mouse model of type one diabetes and examined the interplay among α-, β-, T-cells, myeloid cells, and macrophages in pancreatic islets during the progression of insulitis. Disease evolution was tracked based on T/β cell ratio in individual islets. In the early stage, we found that immune cells are preferentially interacting with α-cell-rich regions of an islet. At the islet periphery α-linked β-cells were found to be targeted significantly more compared to those without α-cell neighbors. Additionally, network analysis revealed increased T-myeloid, and T-macrophage interactions with all β-cells.

## 1 Introduction

### Tracking evolution of islet inflammation

Type 1 diabetes mellitus (T1D) is an autoimmune disorder, characterized by a progressive loss of pancreatic β-cells, decrease in insulin secretion, and an increase in blood glucose levels (Coppieters et al., 2012). T1D islet inflammation, known as insulitis, is associated with β-cell dysfunction and death, has a transient nature in that it clears following complete β-cell loss in an islet, and is usually observed within one year of diabetes onset in humans. Additionally, our work shows that the pathogenic behavior of islet-infiltrating T cells is dependent on the severity of insulitis (Friedman et al., 2014; Lindsay et al., 2015). Previous studies indicate a heterogenous pattern of CD8 T cell infiltration in human pancreatic islets during the early stages of T1D (Coppieters et al., 2012; Rodriguez-Calvo et al., 2015; Damond et al., 2019). Here we utilized the non-obese diabetic (NOD) mouse model - one of the most widely studied diabetic mouse models, with features similar to human T1D (Kikutani and Makino, 1992; Anderson and Bluestone, 2005; Chen et al., 2020). We applied an islet sorting algorithm based on T/β cell ratio in individual islets to account for differences in the amount of infiltration in various islets, that allows to assign a disease-stage to each islet. Islet-wise classification has been previously performed to indicate the stage of insulitis (Carrero et al., 2013; Katsarou et al., 2017; Damond et al., 2019).

### Detangling multi-cellular interactions to understand non-uniform inflammation of an islet

In an islet, peri-insulitis is often observed on one side of an islet (Morgan et al., 2014). We, therefore, asked a question: What is unique about the regions of an islet that are affected by peri-insulitis? With this in mind we then aimed to detangle multi-cellular immune-endocrine interactions by applying network theory analysis – a mathematical method for study relationships between a set of nodes (cells in our case). Network analysis has been successfully applied to analyze protein-protein interactions, gene regulation, and metabolic pathways at the cellular level (Jeong et al., 2000; Barabási and Oltvai, 2004; Schlitt and Brazma, 2007). It has also been used to better understand neuronal architecture and human disease (Bullmore and Sporns, 2009; Barabási et al., 2011). However, more recently it has been applied to conduct *functional* analysis based on coordination of calcium dynamics in endocrine pancreatic islets (Stožer et al., 2013; Johnston et al., 2016; Kravets et al., 2022; Stožer et al., 2022). *Spatial* network analysis is based on the cell-cell proximity and has been previously used to study the organization and interactions of α-, β- and δ-cells in an islet (Hoang et al., 2014; Gosak et al., 2022). Immune cells interact with islet cells through contact-dependent mechanisms, enabling the quantitative analysis of these interactions by assessing the proximity between these cell types (Cnop et al., 2005; Burrack et al., 2017). Specifically, T-cells interact with β-cells via well-defined ligand-based interactions: the T-cell receptor (TCR) on CD8+ T-cells recognizes antigens presented by HLA class I molecules on β-cells, and the Fas ligand on CD8+ T-cells binds to the Fas receptor on β-cells, providing a secondary interaction pathway (Daniels and Jameson, 2000; Dudek et al., 2006; Mariuzza et al., 2020). Additionally, myeloid cells and macrophages interact with β-cells by phagocytosing intact or apoptotic β-cells and their debris, followed by antigen presentation and upregulation of MHC class I and II molecules, recognized by CD8+ and CD4+ T-cells’ TCRs, facilitating further contact-based interactions (Friedl and Gunzer, 2001; Vomund et al., 2015). Islet cells also exhibit paracrine signaling, intensified by closer proximity among cells, which is pivotal in the islet’s functional dynamics (Caicedo, 2013). Recently, spatial network analysis was adopted to show that α-cells are not randomly organized in C57BL/6 mouse islets (Tran Thi Nhu et al., 2017). The number of direct α–α connections was shown to be greater in the experimentally-derived network than a network of randomly connected islet cells (Tran Thi Nhu et al., 2017). In the present study, we utilized spatial networks, where links were assigned based on the cell-cell proximity. To assess whether our observed spatial interactions are merely due to chance, we created random networks by assigning random positions to immune cells, while preserving their empirically derived average distances to the neighboring endocrine cells, and leaving positions of the endocrine cells intact. We then tracked the network evolution over the disease progression pseudo time, based on the T/β spectrum, described above.

Pancreatic tissue slices from non-diabetic NOD mice of ages [16-23 weeks] were used to develop a workflow to quantify heterotypic interactions between α-cells, different β-cell subpopulations (α-linked and non-α-linked), and various immune cell types. We sought to investigate if β-cells closer to α-cells, interact more with immune cells during T1D, as the destruction of these cells could explain the loss in the first phase of insulin response observed in individuals with pre-diabetes.

## 2 Materials and Methods

### Animal care

NOD/ShiLtJ mice (001976) were obtained from The Jackson Laboratory and bred in-house. All animal procedures were approved by the Institutional Animal Care and Use Committee at the University of Colorado Anschutz Medical Campus.

### Immunofluorescence and imaging of mouse pancreas

WT NOD females were 16-23 weeks of age, with blood glucose readings below 250mg/dL. The whole pancreas was excised and fixed in formalin (VWR) for 24 hours. After fixation, samples were stored in 70% ethanol until paraffin embedding. 4-µm sections were cut. Embedding and sectioning were performed by the University of Colorado Anschutz histology core. Samples were stained using a Leica Auotstainer and imaged using whole tissue scanning on the Akoya Polaris Imaging System.

In collaboration with the Human Immune Monitoring Shared Resource (HIMSR) at the University of Colorado School of Medicine, we performed 7 color multispectral imaging using the Akoya Biosciences Vectra Polaris instrument. Six markers were utilized, including insulin (β-cells), glucagon (α-cells), CD11c (for myeloid cells, including dendritic cells, monocytes, and macrophages), CD3 (for T cells), F4/80 (for macrophages). Myeloid cells include any cell type that develops/differentiates from a common myeloid progenitor. Myeloid cells include two major categories: (1) mononuclear phagocytes and (2) granulocytes. This experiment stains for mononuclear phagocytes, which are comprised of 3 myeloid subsets: Monocytes, Macrophages, Dendritic Cells. This experiment does not stain for granulocytes (Neutrophils, Eosinophils, Basophils, Mast cells), which are also myeloid subsets.

The slides were stained on the Leica Bond RX autostainer according to standard protocols provided by Leica and Akoya Biosciences. Briefly, the slides were deparaffinized, heat treated in antigen retrieval buffer, blocked, and incubated with primary antibody, followed by horseradish peroxidase (HRP)-conjugated secondary antibody polymer (Akoya), and HRP-reactive OPAL fluorescent reagents (Akoya) that use TSA chemistry to deposit dyes on the tissue immediately surrounding each HRP molecule. To prevent further deposition of fluorescent dyes in subsequent staining steps, the slides were stripped in between each stain with heat treatment in antigen retrieval buffer (Akoya). Whole slide scans were collected by widefield fluorescent imaging using a 20x/NA = 0.75 objective and a pixel size of 0.5mm/px. inForm software was used to unmix the 7 color images and subtract autofluorescence. Table 1 describes the reagents, including antibodies and their specific details such as suppliers, catalog numbers, clones, OPAL dye wavelengths, and antigen retrieval conditions used in immunofluorescence staining protocols.

**Table 1.** Summary of immunofluorescence staining reagents used in our study.

| Target | Supplier | Catalog number | Clone | OPAL Dye | Antigen retrieval |
| --- | --- | --- | --- | --- | --- |
| Ms CD3 | Cell signaling technology | 999440S | D4V8L | 570 | pH9 |
| Ms Insulin | Cell signaling technology | 4590S | Polyclonal | 480 | pH6 |
| Ms CD11c | Cell signaling technology | 97585S | D1V9Y | 520 | pH6 |
| Ms Glucagon | Cell signaling technology | 2760S | Polyclonal | 620 | pH6 |
| Ms F4/80 | Cell signaling technology | 30325S | D4C8V | 780 | pH6 |

### Image analysis

All image analysis was primarily done in ImageJ (FIJI) using pre-existing plugins (Schindelin et al., 2012). Akoya Polaris whole slide viewer software (Akoya Biosciences and The Spatial Biology Company, Marlborough, United States of America), and Imaris (Bitplane, Zurich, Switzerland) were used for nuclei segmentation.

### Pre-processing

A smaller section, corresponding to a single islet was first manually segmented from a whole slide image, that represented the entire pancreatic slice. To account for bleed through of signal from the insulin channel into the other channels, we subtracted the regions corresponding to β-cells detected in the insulin channel from the CD11c, CD3 and F4/80 channels. As the β-cells in the pancreas are not anticipated to exhibit positive staining for the immune cell markers, we reasoned that the observed immune cell marker signal in regions of β-cells is likely due to bleed through. As expected, we observed highest amount of bleed through between the spectrally-neighboring insulin and CD11c channels (see also Colocalization Analysis section below). Figures S3C and S3D represent the CD11c channel of an islet before and after bleed through correction. The outline of the islet was manually sketched around the α- and β-cells. The islet selection was then enlarged by 60 μm to encompass the immune cells present in the periphery of the islet. The selection was then replicated in the images corresponding to other channels. Finally, the images were filtered by a median filter of radius 2 pixels to minimize background noise.

### Nuclei Segmentation

For every islet, the channel corresponding to its DAPI staining was overlaid with another channel of interest in the Akoya Polaris whole slide viewer software to visualize the locations of the nuclei of the corresponding cell type. The image corresponding to that channel was loaded in Imaris. Spots were manually added in Imaris by referring to the overlaid image in the whole slide viewer software to represent the nuclei. The positions of the nuclei (as X and Y coordinates) were then exported from Imaris for further analysis.

### Colocalization Analysis

To test colocalization of the signal from different channels in the same cell, we calculated the Manders coefficients using the previously reported colocalization threshold plugin in ImageJ (Manders et al., 1993; Costes et al., 2004). The Manders colocalization coefficients between two channels - 1 and 2 of ‘n’ pixels of intensities ‘P1’ and ‘P2’ can be approximated as shown in equations (1) and (2) (Manders et al., 1993; Costes et al., 2004): -

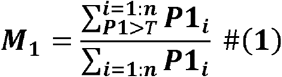

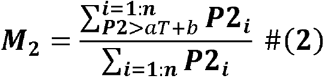

Here, ‘T’ is an intensity threshold for channel 1 that was automatically determined by the plugin’s iterative algorithm, and ‘aT + b’ is its corresponding intensity in channel 2 (constants ‘a’ and ‘b’ were determined by fitting the intensities of channel 1 with channel 2). Threshold ‘T’ was first initialized with a high value of intensity. Pearson’s correlation coefficient was computed between the channels for the pixels whose intensities are below threshold ‘T’ in channel 1 and ‘aT + b’ in channel 2. Threshold ‘T’ was decreased until this correlation coefficient was zero. Results are shown in Supplementary Figure S1.

### Ranking of insulitis progression

Progression of insulitis was ranked by computing the ratio of T-cells to β-cells (insulitis degree), as shown in equation (3). For every islet, the number of T-cells and β-cells that were inside the islet plus within 60 μm of the islet rim were considered, and their ratio was computed using equation 3. We then categorized the islets as early-, intermediate- and late-stage islets on a spectrum. We categorized the first 44 islets in this spectrum with T/β cell ratios between 0 and 0.51 as early-stage insulitis, the next 46 islets with T/β cell ratios between 0.51 and 1.25 as intermediate-stage insulitis, and the last 44 islets with T/β cell ratios greater than 1.25 as late-stage insulitis.

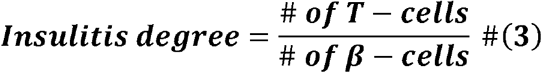

As detailed in results, we observed heterogeneity in the amount of peri-insulitis and infiltration of different islets in the same mouse (see Supplementary Figure S4F for mouse by mouse distribution of insulitis degrees). Mouse #5 had 14 islets with T-cells to β-cells ratios ranging from 0.065 to 5.692. Six of these islets were classified as early-stage insulitis, 3 islets were classified as intermediate stage insulitis, and 5 islets were classified as late-stage insulitis. Similarly, mouse #2 had 1 early-stage islet, 4 intermediate-stage islets and 6 late-stage islets with ratios ranging from 0.257 to 7.800. Different stages of islet inflammation are shown in Figure 2A.

### Cellular network construction

Cellular interactions were modelled as a graph, where cells were considered as nodes and links were assigned between the nodes based on their proximity to each-other. The islet mask was enlarged by 20 μm, and all immune cells located within this mask were considered to be in the network. Since, we referred to the position of a cell using the position of its nucleus, all distances specified in this paper correspond to center-to-center distances between cells. The distance for cells to be considered interacting varied between 15 μm, 20 μm and 25 μm (results for each option were reported separately). Additionally, links were constructed only between cells of differing cell types, as we are interested only in the heterotypic interactions between cells. Two-cell networks were constructed between α- and β- cells, α- and T- cells, α-cells and macrophages, α- and myeloid cells, and three-cell networks were constructed between [α-, β-, T-] cells, [α-, β-, macrophage] cells and [α-, β-, myeloid] cells for further analysis.

### Network analysis

All network construction and network analysis algorithms were done in MATLAB using custom scripts. The amount of cellular interactions between two cell types were quantified by computing the average number of connections, links (K_avg_) of a particular cell type in a two-cell network. Average links refers to the mean number of connections a cell has in the network. It can be mathematically expressed as shown in equations (4) and (5). Three-cell networks were used to quantify the interactions between T-cells/macrophages/myeloid cells and α-linked β- cells vs non-α-linked β- cells. Specifically, β-cells linked to α-cells were first identified, and then two-cell networks were created between α-linked β- cells and one immune cell type. Same was done for the non-α-linked β- cells and one immune cell type.

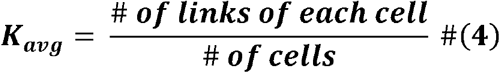

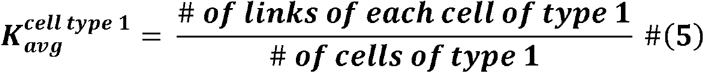

### Randomizing procedure

This randomization procedure was used to create random heterotypic networks for assessment of likelihood of our observations being due to chance. The positions of the α- and β-cells were fixed, and the immune cells were randomly distributed around the islet rim. The immune cells were classified into two categories – inside the islet and outside the islet, based on their location with respect to the islet rim. First, the immune cells outside the islet were considered, and the distance of each cell to the islet rim was computed. A normal distribution of 1000 distances was created using the mean and standard deviation of these experimentally-determined distances. Next, the algorithm randomly replaced each immune cell’s distance with a random distance from the above-mentioned normal distribution. In this manner, each immune cell was randomly repositioned withing the islet periphery, while average distance of all immune cells to the islet rim almost remained the same. The same procedure was adopted to randomize the positions of immune cells inside the islet. The randomization process was repeated 100 times. Each time (each seed), the set of distances randomly picked from the normal distribution, and the positions randomly assigned to cells, were varied.

### Outer islet cell analysis

In order to account for 1) α-cells being predominantly located at the islet periphery, and b) islet cells at the islet periphery having more opportunity to interact with the immune peri-insulitis, compared to cells located in the islet core, we performed analysis of the *outer cell layer*, as described further. Only cells at a distance 20 μm from the islet rim (20 μm inside the islet and 20 μm outside the islet) were considered. Three-cell networks were created using i) α-, (ii) β- cells in the outer layer, and (iii) each of the immune cell types. β-cells linked to α cells were classified as “α-linked β-cells”, and all other β-cells in the outer ring were classified as non-α-linked. The number of links between α-linked β-cells and immune cells were computed and compared to the number of links between non-α-linked β-cells and immune cells.

### Determination of islet and immune cell proportions in islet halves

To assess whether immune cells preferentially infiltrate α-cell rich regions within an islet, we divided the islet mask into two halves along a vertical or horizontal plane, as illustrated in Supplementary Figure S5A. We then calculated the number of islet cells (α- or β-) and immune cells (T-, macrophage, or myeloid cells) in each half, and then computed the proportion of each cell type within each section. **Network analysis to study inter-islet variations in islet size**

To explore whether inter-islet size differences affect our observed trends, we categorized islets into three groups based on their cross-sectional area: small (0 – 7000 μm^2^), medium (7000 – 20000 μm^2^), and large (>20000 μm^2^). Each group was then subdivided into early, intermediate, and late-stage insulitis based on their insulitis degrees. We then repeated various network analyses for each group, as shown in Supplementary Figures S10 and S12.

### Statistical analysis

All statistical analysis was performed using GraphPad Prism (GraphPad, Boston, United States of America). Data was reported as ± SEM. The differences were considered to be statistically significant for p < 0.05. The tests used for evaluating statistical significance between results are mentioned in the figure captions.

## 3 Results

### Network analysis can be successfully used as a tool to quantify cellular interactions

We first sought to develop a metric that could quantify cellular interactions using the positions of various cells in an islet. Pancreatic tissue slices were stained for DAPI, insulin, glucagon, CD3, CD11c and F4/80 and whole slide images were captured. As depicted in Figure 1A and 1B, an islet was first cropped from a whole slide image of the cross section of a mouse’s pancreas. Nuclei of cells were segmented for each of the channels, and cross-referenced with the DAPI channel (Figure 1C-G). Euclidean distances were then computed between the cells, and finally, proximity-based networks were created using a suitable threshold as shown in Figure 1H (See Materials and Methods section for detailed description). We only considered the immune cells that were within 20 μm of the islet edge, and all immune cells inside the islet. For network link assignment we tried different thresholds - 15, 20 and 25 μm, and saw same trends (more or less pronounced based on the threshold) in our findings for each of them (Supplementary Figures S3, S8, S11). Therefore, we chose to use 20 μm as the threshold for network analysis. Finally, we observed modest colocalization between the immune cell markers (see Supplementary Figure S1 for more information), and, after image pre-processing, could ascertain that the CD3 channel correspond to T cells, CD11c to myeloid cells, and F4/80 to macrophages (ML□). We primarily sought to study the heterotypic interaction between two cell types, and we therefore generated networks for two types of cells at a time. We analyzed a total of 134 islets from 11 different mice (See Supplementary Figure S2 for more information on the age, glucose recordings and the number of islets analyzed for each mouse).

**Figure 1.**
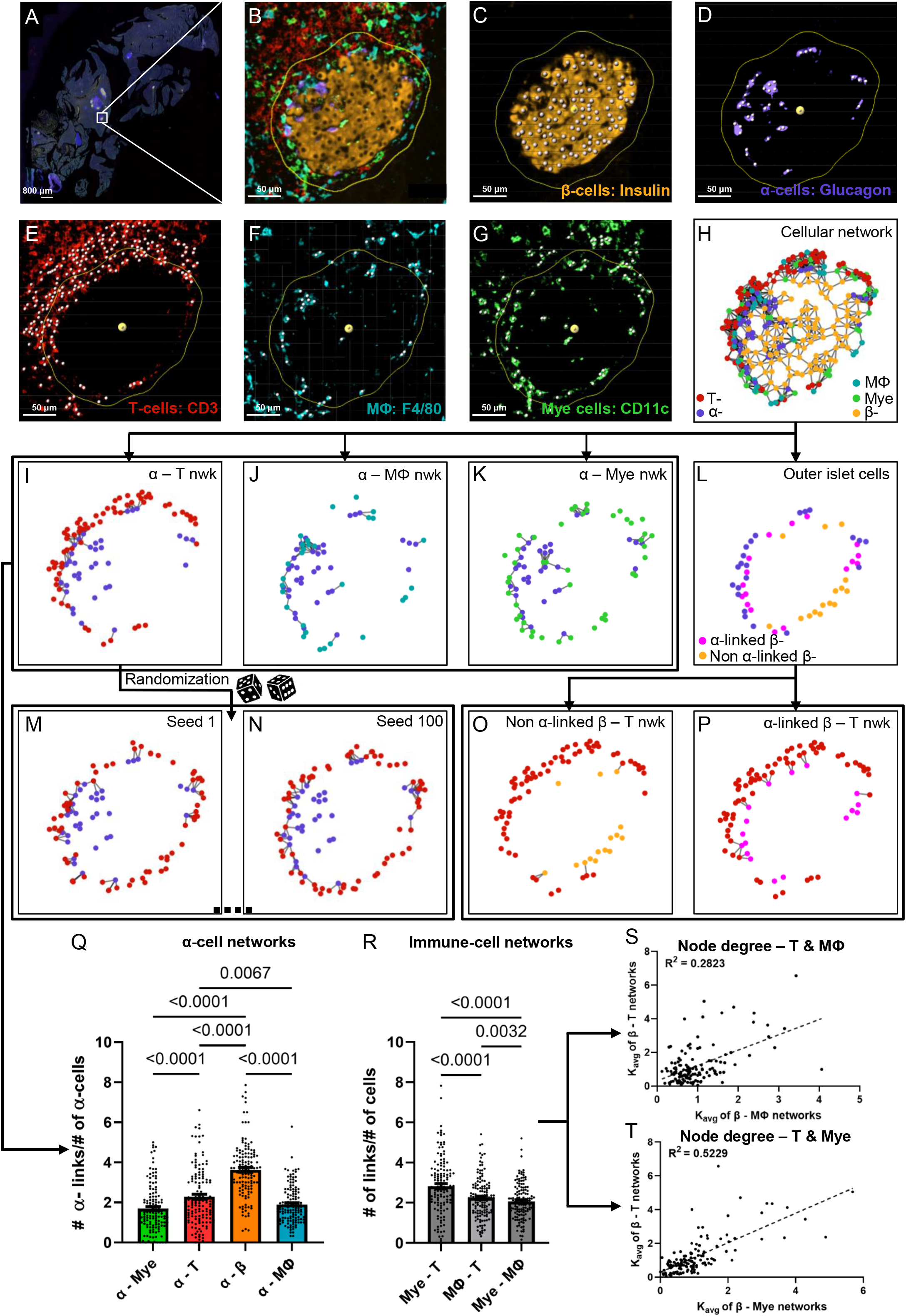
**(A)** Representative whole slide image of a pancreatic slice, white box represents the islet of interest. **(B)** Representative islet cropped from the whole slide image shown in A. **(C)** β-cell nuclei segmented from the insulin signal extracted from the islet shown in B. **(D)** α-cell nuclei segmented from the glucagon signal extracted from the islet shown in B. **(E)** T-cell nuclei segmented from the CD3 signal extracted from the islet shown in B. **(F)** Macrophage (MLJ) nuclei segmented from the F4/80 signal extracted from the islet shown in B. **(G)** Myeloid (Mye) cell nuclei segmented from the CD11c signal extracted from the islet shown in B. Yellow outlines in B - G represents distance of 20 μm from the islet rim, white spots in C -G represents nuclei. **(H)** Representative network of the islet shown in B created using the cellular positions obtained in C – G. **(I) – (K)** represent α-T, α-macrophage and α-myeloid cell networks of the islet shown in B created using cellular positions obtained in C - G. **(L)** Representation of the outer layer of islet cells of the islet shown in B created from the network shown in H. **(M)** and **(N)** represent random networks obtained by randomizing the positions of T-cells in the network shown in I. **(O)** and **(P)** represent the α-linked β-T and non α-linked β-T networks created using the outer layer of islet cells obtained in L and the positions of T-cells obtained in E. **(Q)** Comparison of K_avg_ of different α**-**cell networks using network analysis (n = 134 islets from 11 mice) (RM one way ANOVA with Geisser-Greenhouse correction and Tukey’s multiple comparisons tests were used for statistical analysis) **(R)** Comparison of K_avg_ of different immune cell networks using network analysis (n = 134 islets from 11 mice) (RM one-way ANOVA with Geisser-Greenhouse correction and Tukey’s multiple comparisons tests were used for statistical analysis). **(S)** Relationship between the node degrees of β-macrophage and β-T networks obtained using network analysis (n = 134 islets from 11 mice) (Simple linear regression was used for statistical analysis). **(T)** Relationship between the node degrees of β-myeloid and β-T networks obtained using network analysis (n = 134 islets from 11 mice) (Simple linear regression was used for statistical analysis). See Materials and Methods for detailed description on pre-processing, segmentation, network analysis, outer-islet cell analysis and randomization.

As a first proof of concept, we compared the average number of connections (links), K_avg_ of α-cells in α – immune cell networks, and α – β cell network and observed the highest K_avg_ in α – β cell network, which suggests that α-cells interact the most with β-cells, as expected (see Figure 1Q). Figures 1I-K represent the various α – immune networks for the islet shown in Figure 1B. Secondly, as a proof of concept we compared the interactions between the different immune cell types cumulative for all islets by comparing the average links, also referenced through the text as *node degrees* or K_avg_ between T – myeloid cell networks, T cell – macrophage networks and myeloid cell – macrophage networks (see Figure 1R). T - myeloid cell networks had a higher and significantly different K_avg_ compared to other immune cell pairs. These findings reinforce previous studies which state that T cells and myeloid cells interact with each other during T1D (Calderon et al., 2011; Sandor et al., 2019; Jain et al., 2020), and support our use of the network analysis for assessment of cellular interaction. Lastly, to assess which immune cell *pairs* interact stronger with β-cells, we plotted the node degrees of various β – immune cell network pairs against each other and observed that β – myeloid cell network had the highest correlation coefficient (R^2^) with the β – T cell network (see Figure 1S,T and Supplementary Figure S3E). This suggests that interaction of T-cells with β-cells correlates strongly with myeloid cell proximity and vice versa.

### α-cell-rich regions of an islet are more infiltrated and characterized by non-random T- α and macrophage-α cell interactions

To track evolution of multicellular networks with peri-insulitis and insulitis progression, we computed the ratio of the number of T-cells and β-cells in each islet and sorted them in ascending order in Figure 2A (see detailed description in Materials and Methods). T–β cell ratios ranged from 0.0217 to 27.50. T–β cell ratios of islets of different pseudo-stages/stages of insulitis were compared in Figure 2B. The degree of infiltration in our early-, intermediate-, late-stage islets resembled the islets that were previously categorized as mild, moderate, and severe insulitis (Miyazaki et al., 1985). As a proof of concept, we also plotted the T-cell density of the islets in the same spectrum obtained by sorting the ratios (psuedotime) and observed a similar exponential pattern as shown in Supplementary Figure S4A. The exponential patterns observed where also fit using log scales as shown in Supplementary Figures S4B and S4D. Next, we plotted the correlation between the number of β-cells and T-cells in an islet (Supplementary Figure S4E). By color-coding the stages, we observed distinct clusters with varying slopes, aligning within the predefined insulitis degree ranges for each stage.

**Figure 2.**
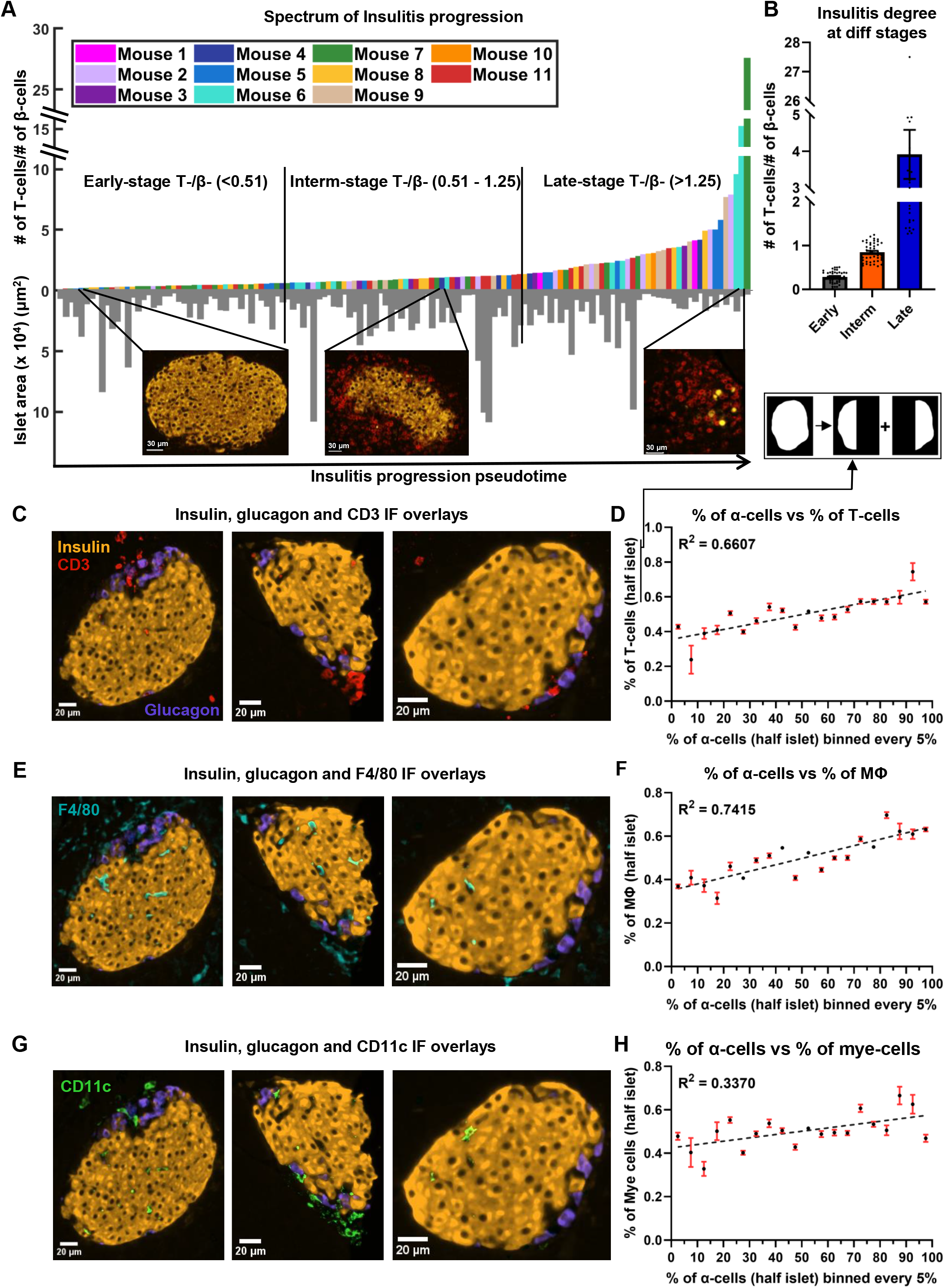
**(A)** Progression of insulitis in different islets, sorted using T/β cell ratios (psuedotime) (n = 134 islets from 11 mice). Orange and red colors in the immunofluorescent images correspond to insulin and CD3 stains. **(B)** Comparison of insulitis degrees of islets classified as early- (n = 44 islets), intermediate- (n = 46 islets) and late- (n = 44 islets) stage insulitis. **(C)** Immunofluorescence (IF) overlay of insulin (orange color), glucagon (purple color), and CD3 (red color) stains in three early-stage representative islets. **(D)** Relationship between the proportion of α-cells and the proportion of T-cells in half islets at a distance of at a distance of 20 μm from the islet rim (n = 134 islets). **(E)** IF overlay of insulin, glucagon, and F4/80 (cyan color) stains in three early-stage representative islets. **(F)** Relationship between the proportion of α-cells and the proportion of macrophages in half islets at a distance of at a distance of 20 μm from the islet rim (n = 134 islets). **(G)** IF overlay of insulin, glucagon, and CD11c (green color) stains in three early-stage representative islets. **(H)** Relationship between the proportion of α-cells and the proportion of myeloid cells in half islets at a distance of at a distance of 20 μm from the islet rim (n = 134 islets). See Materials and Methods for detailed description on the algorithm used to sort the islets and for how the proportion of α-cells and immune in half islets were determined. Simple linear regression was used for statistical analysis in D, F and H.

#### Infiltration spectrum

As shown in the inserts in Figure 2A, islets at the beginning of the spectrum had sparse T-cells and intact β-cell mass (mostly peri-insulitis), islets at the center of the spectrum had an appreciable amount of intact β-cells and an increase in the number of T-cells (insulitis), and finally islets at the end of the spectrum were destroyed, and had significant β-cell loss (destructive insulitis). In Supplementary Figure 4F we demonstrate the observed high heterogeneity in T-cell presence of different islets in the same mouse. The heterogeneity between mice in terms of T-cell presence has been previously observed (Johansson et al., 2003). On the other hand, the variability of islet inflammation in the *same* mouse been known but often hindered by reporting degree of insulitis in a mouse-averaged manner. Recently, categorizing insulitis progression (i.e., identifying a pseudo stage of the disease) in the islet-based manner have been used more widely (Carrero et al., 2013; Friedman et al., 2014; Lindsay et al., 2015; Katsarou et al., 2017; Damond et al., 2019).

#### T cells and macrophages non-randomly interact with a-cells

We next sought to determine which immune cell type interacted the most with α-cells during the early stages of islet inflammation. We found that α – T cell networks had a higher number of links compared to α – macrophage and α – myeloid networks (p = 0.0067 and p < 0.0001) (see Figure 1Q). To determine whether the observed interactions between the immune cells and α-cells are due to chance, or if immune cells were drawn to α-cells, we generated random (or shuffled) networks where we fixed the positions of α-cells (same as experimentally observed) and randomly shuffled the positions of immune cells as shown in Figures 1I, 1M and 1N (see Materials and Methods and Supplementary Figure S6 for detailed description). After randomization, we observe a decrease in the number of α - immune cell links (Figure 3). An example of such non-uniform and non-random distribution of the immune cells is shown in the representative early-stage islet in Figures 3A, 3C and 3E. The upper region of the islet had more α-cells than the lower region. Perhaps as a result of that, the number of immune cells in the upper region of the islet was also higher. In the random (simulated) networks there was no significant difference of immune cell numbers in any regions of the islet.

**Figure 3.**
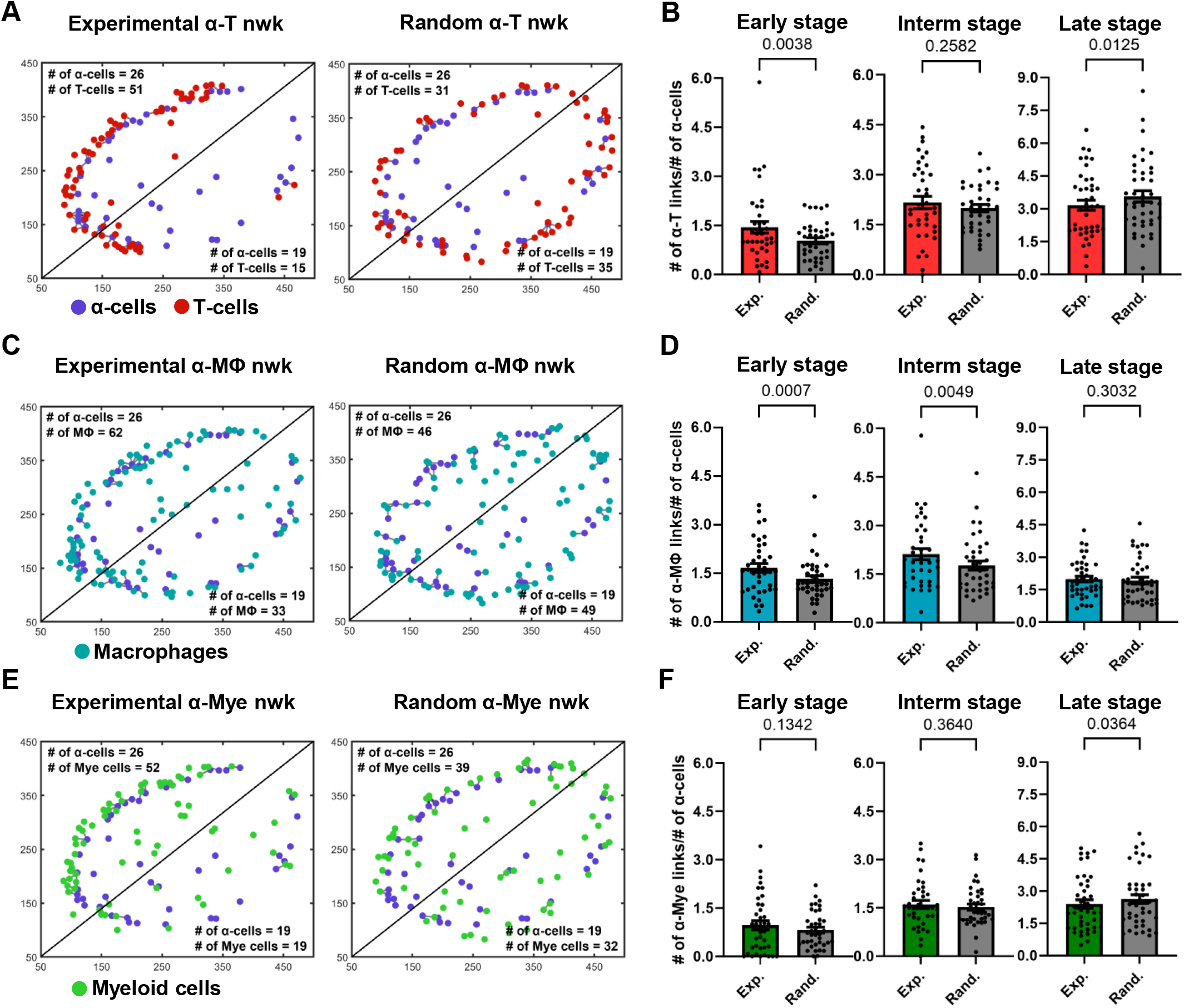
**(A)** Experimental and random α-cell – T-cell networks of a representative early-stage islet. **(B)** Comparison of K_avg_ of experimental and random α-cell – T-cell networks (nwks) in early (n = 39 islets), intermediate (interm) (n = 38 islets), and late (n = 42 islets) stages using network analysis. **(C)** Experimental and random α-cell – macrophage (MLJ) networks of a representative early-stage islet. **(D)** Comparison of K_avg_ of experimental and random α-cell – macrophage networks in early (n = 39 islets), intermediate (n = 38 islets), and late (n = 42 islets) stages using network analysis. **(E)** Experimental and random α-cell – myeloid (mye) cell networks of a representative early-stage islet. **(F)** Comparison of K_avg_ of experimental and random α-cell – myeloid cell networks in early (n = 39 islets), intermediate (n = 38 islets), and late (n = 42 islets) stages using network analysis. Paired parametric t tests were used for statistical analysis in B, D and F. See Materials and Methods for detailed description on randomization and network analysis.

#### α-immune interactions become random as disease progresses

We then computed the average number of links of these random networks and compared them with the average number of links, K_avg_, of the experimental networks in early-, intermediate- and late-stages of the disease. In Figures 3B and 3D, we show that in the early stages of the disease the K_avg_ of experimental α – T cell and α – macrophage networks was greater and more statistically significant than the corresponding random networks (p = 0.00380 and p = 0.0007). Experimental α – macrophage networks also exhibited more interactions than random networks in the intermediate stages (p = 0.0049). However, in the intermediate and later stage α – T cell networks the difference was not statistically significant and no significant differences were observed in the later stages of α – macrophage networks. These findings suggest that in the early stages of the disease the α-T as well as α-macrophage interactions, are not due to a chance. While we also observed an increase in the number of myeloid cells in α-cell-rich regions of an islet (Figure 3E), the statistical significance was not observed between experimental and random α – myeloid cell networks (Figure 3F). Additionally, we observed similar results when we fixed the positions of immune cells and shuffled the positions of α-cells (see Supplementary Figure S7).

### α-linked β-cells interact with T cells and macrophages stronger than non- α-linked

In most of the islets classified to have an early-stage inflammation (examples are in Figures 2C, 2E, and 2G) we observed that the α-cell-rich regions of the islet had more immune cells. This trend was diminished in the later stages of the disease. One can see more examples of this pattern in the Figure S9 where 7 out of 10 randomly picked early-stage islets exhibited this pattern of infiltration. We divided each islet into two regions using a vertical plane and observed a positive correlation between the percentage of α-cells and the percentage of immune cells in each region. (Figures 2D, 2F, and 2H) We similarly noted a trend when employing a horizontal plane (Supplementary Figure S5B). Plotting the relationship between the percentage of β-cells and the percentage of immune cells revealed no correlation (see Supplementary Figures S5C and S5D), further indicating immune cells’ preference for infiltrating α-cell rich regions of an islet. In order to de-tangle whether this effect is due to α-cells themselves, or α-linked β-cells, we then sought to quantify whether the β-cells closer to α-cells interact more with immune cells. To compensate for the fact that β-cells in the islet core have less chances to interact with immune cells at the early stages of insulitis, *i*.*e*., to provide equal opportunity for both α-linked β-cells and non-α-linked β-cells to interact with immune cells near islet’s surface, we considered only the peripheral (outer ring) β-cells (see Figure 4A and 4B) (see materials and methods for detailed description on how the outer ring was identified). Withing the outer rings we constructed networks between α-linked β-cells and immune cells, and non-α-linked β-cells and immune cells (see Figure 1L, 1O, 1P). We compared the K_avg_ of these networks at different stages of inflammation and found that the K_avg_ of α-linked β – T cell network was significantly greater than the K_avg_ of non-α-linked β – T cell networks in the early (p < 0.0001) stage (Figure 4D). Similarly, the K_avg_ of α-linked β-cell – macrophage networks was significantly greater than the K_avg_ of non-α-linked β-cell – macrophage networks in the early (p = 0.0010) and intermediate (p = 0.0053) stages (Figure 4F). However, in the later stages the differences were no longer significant. No statistically significant trends were observed comparing the K_avg_ of α-linked β – myeloid cell and non-α-linked β – myeloid cell networks (Figure 4F). These observations suggest that the β-cells situated next to the α-cells (or interacting with α-cells) might interact more with the immune cells in the earlier stages of the disease. This is a novel finding not previously reported elsewhere.

**Figure 4.**
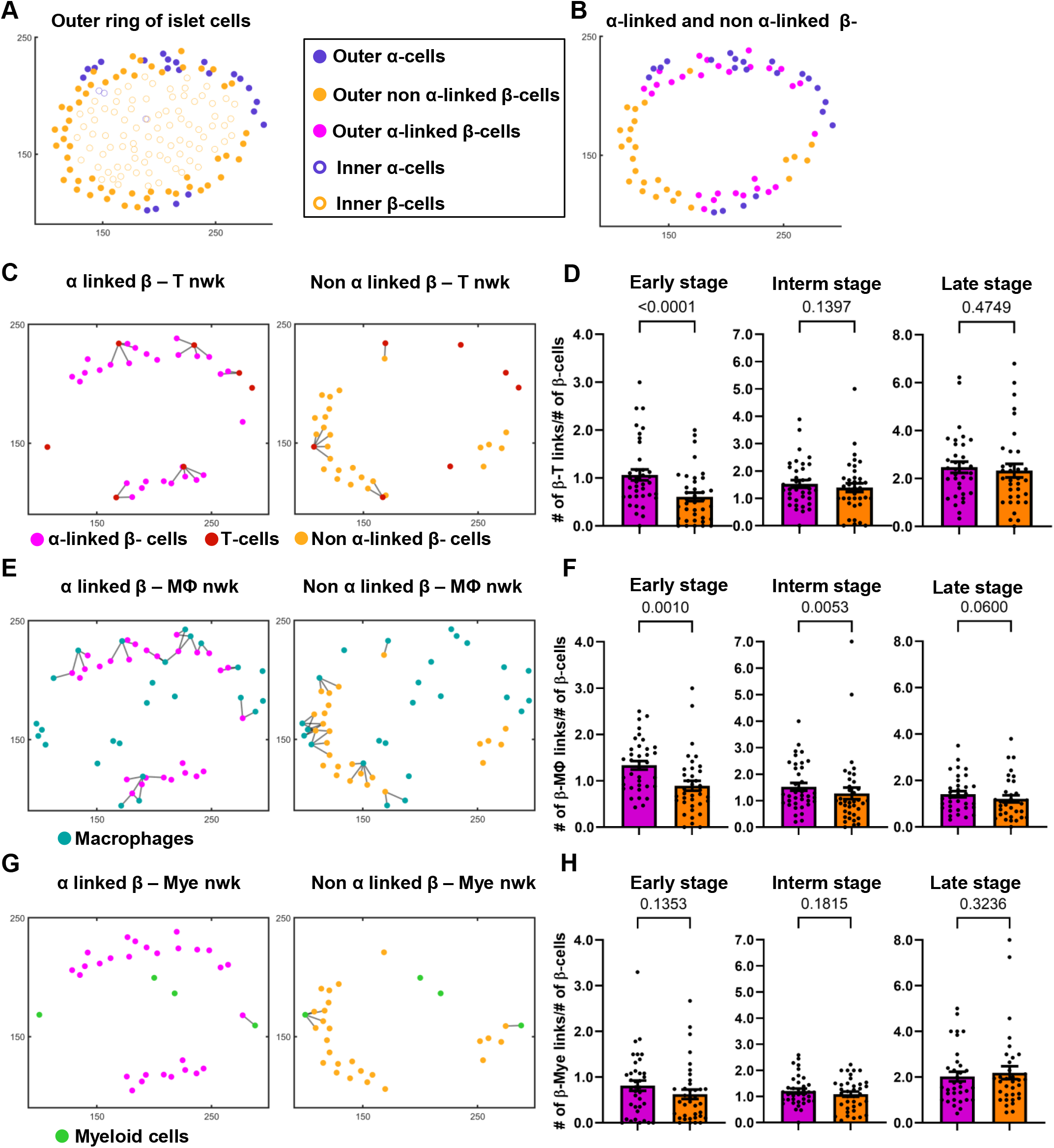
**(A)** Differentiation between the outer ring and inner core of islet cells, of a representative islet. **(B)** Identification of α-linked and non α-linked β-cells in the outer ring of islet cells. **(C)** α-linked β-cell – and non-α-linked β-cell – T-cell networks (nwks) of a representative early-stage islet. **(D)** Comparison of K_avg_ of α-linked β-cell - and non-α-linked β-cell – T-cell networks in early (n = 37 islets), intermediate (interm) (n = 37 islets), and late (n = 34 islets) stages using network analysis. **(E)** α-linked β-cell – and non-α-linked β-cell – macrophage (MLJ) networks of a representative early-stage islet. **(F)** Comparison of K_avg_ of α-linked β-cell and non-α-linked β-cell – macrophage networks in early (n = 37 islets), intermediate (n = 37 islets), and late (n = 34 islets) stages using network analysis. **(G)** α-linked β-cell – and non-α-linked β-cell – myeloid (mye) cell networks of a representative early-stage islet. **(H)** Comparison of K_avg_ of α-linked β-cell and non-α-linked β-cell – myeloid cell networks in early (n = 37 islets), intermediate (n = 37 islets), and late (n = 34 islets) stages using network analysis. Wilcoxon matched-pairs signed rank test was used for statistical analysis. See Materials and Methods for detailed description on network analysis and on how the outer ring of islet cells were isolated.

## 4 Discussion

We successfully designed a workflow that could be used to quantify the cellular interactions using the proximity of cells in pancreatic islets during insulitis progression. In our dataset, we observed presence of T-cells both in the periphery of the islet and inside the islet. Previously it was shown that the T-cells could traffic through the islet vasculature and extravasate into the islet (Sandor et al., 2019), explaining presence of the T cells inside the islet. In the early stage we observed majority of the immune cell outside the islet (peri-insulitis). Our analysis showed β-cell associations with myeloid-T-cells pairs, and macrophages-T-cell pairs. These observations can be rationalized by myeloid cells and macrophages being antigen presenting cells, that release various chemokines responsible for T-cell recruitment and antigenic stimulation (Friedman et al., 2014; Lindsay et al., 2015; Sandor et al., 2019).

### Why α-cell-rich areas of an islet have more leucocytes

In human islets, α-cells were reported to express more IL-1β, IL-6, and other non-classical MHC class 1 molecules than β-cells during T1D (Anquetil et al., 2017; Benkahla et al., 2020; Rajendran et al., 2020a; Rajendran et al., 2020b). Higher expression of pro-inflammatory cytokines, could increase inflammation in the α-cell-rich regions of the islet, and potentially explain why immune cells are drawn to these regions in the early stages of disease. In 12 to 21 week old NOD mice, α-cells were reported to express more CXCL10 (a chemokine recognized by the CXCR3 receptor of T-cells) (Nigi et al., 2020). Glucagon – CXCL10 colocalization, was greater than insulin – CXCL10 colocalization in new onset diabetic NOD mice, and further increases on diabetes onset (Nigi et al., 2020). Increased expression of CXCL10 by α-cells could also potentially explain the presence of more T-cells and macrophages in the α-cell rich regions of an islet (Figures 2C, 2D, 2E and 2F). Previous studies have indicated that the increased expression of CXCL10 increases T-cell and macrophages infiltration in the islet, and that diabetogenic T-cells migrate to sites containing higher levels of CXCL10 (Christen et al., 2004; Tanaka et al., 2009; Roep et al., 2010). It is important to note that while α-cells interact with T-cells and macrophages in the early stages, the α-cells still remain alive until very advanced stages of diabetes (Bru-Tari et al., 2019). Additionally, in the later stages of islet inflammation, we do not notice specific interactions between the α-cells and immune cells (Figures 3B, 3D and 3F), possibly due to an increase in myeloid cell infiltration, myeloid cell – T cell interactions, and an overall increase in CXCL10 expression throughout the islet (Shigihara et al., 2006; Sandor et al., 2019).

### Why α-linked β-cells have more leucocyte interactions

Firstly, β-cell proximity to α-cells increases their exposure to glucagon and GLP1, which is sensed via GLP1-R on the surface of the β-cells and increases cyclic AMP and insulin production (Moens et al., 1998; Marchetti et al., 2012). Our discovery of T-cells and macrophages non-random interaction with α-linked β-cells may be rationalized by higher amounts of insulin produced by α-linked β-cells. This effect is more pronounced in larger sized islets (see Supplementary figure S12), as there are fewer β-cells in smaller-sized islets, leading to a higher proportion of β-cells being influenced by the glucagon released by α-cells. Secondly, in humans and rodents, β-cells exhibit a distinct biphasic insulin response to glucose stimulation (Curry et al., 1968; Porte and Pupo, 1969; Nunemaker et al., 2006). There is a marked loss in the first phase of insulin secretion in individuals with pre-diabetes and with T1D (Brunzell et al., 1976). We previously showed that, first responder β-cells are a subpopulation of β-cells that drive this first phase of islet’s response to glucose, and that ablation of these 1^st^ responder β-cells leads to a loss in first phase response (Kravets et al., 2022). First responder β-cells are also located closer to α-cells (Kravets et al., 2023), which combined with our finding in this work, suggests that it is likely that first responder β-cells are affected sooner than other β-cells during insulitis. This, in turn, means that loss of the first phase of insulin secretion may be due to immune-mediated destruction of the first responder β-cells by the T cells.

### The main limitations of our study are as follows

Our work was done on the fixed tissue, and the networks were constructed based on the proximity of the immune cells to endocrine cells. It would be more beneficial to study dynamic interactions in live tissue, which is relatively difficult to do due to scarcity and fragility of the live human tissue samples. Additionally, we studied cellular interactions in a 2D tissue, studying cellular interactions in a 3D microenvironment with addition of the markers for vasculature will provide more insights into the changes in islet architecture with disease progression. Moreover, the CD3+ T-cells in our study encompass both helper and cytotoxic T-cells, it would be more beneficial to study the interactions between islet cells and each subpopulation of T-cells separately, as the two types of T cells have different roles in the pathogenesis of T1D. Finally, the network analysis technique employed in this study assumes that cells are interacting based on the proximities of the cells, however studying functional networks, *in vitro* and *in vivo* will provide more insights. Future studies can focus on the role of α-cells in the early stages of T1D, and what properties of α-cells, and α-linked β-cells makes them more susceptible to immune attack.

## 5 Conclusion

Overall, the findings indicate dynamic changes in endocrine-immune cell interactions during the progression of insulitis and shine new light on the α-linked β-cells, or α-cells as potential primary targets. Non-random T-cell and macrophage interactions with α-cells, polarization of peri-insulitis and insulitis patterns towards α-rich regions of the islet early in disease progression, and most importantly, significand prevalence of α-linked β-cell interactions with immune cells suggests that these cells will be lost first in T1D.

## 7 Author Contributions

NB performed data pre-processing and analysis, wrote MATLAB scripts, and drafted manuscript; JW designed the immunofluorescence panel, RF generated experimental images of the NOD pancreatic tissue (unpublished previously), proposed the idea of T/β cell spectrum, determined the timing of the tissue collection, and edited the manuscript; LP performed statistical analysis of the data. VK designed hypotheses of the work, quantitative approaches, and edited/wrote the manuscript.

## 9 Acknowledgements

We are grateful for constructive suggestions from Dr. Marko Gosak who provided his feedback at the early stage of this project, Kimberly Jordan and Angela Minic in the Human Immune Shared Resource, and Brittany Basta for support of our animal colony.

## 10 Data Availability Statement

*The dataset reported in this study will be uploaded into the data repository after the manuscript acceptance*.

## 14 Supplemental figure legends

**Supplementary Figure S1**. Mander’s colocalization coefficients (M1 and M2) of the immune cell markers (CD3, CD11c and F4/80) utilized in our study.

**Supplementary Figure S2.** Information on age, blood glucose recordings, number of islets analyzed and the range of insulitis degrees of all the mice utilized in our study,

**Supplementary Figure S3. (A)** Comparison of Kavg of different α-cell networks (created using different thresholds) using network analysis (n = 134 islets from 11 mice) (RM one way ANOVA with Geisser-Greenhouse correction and Tukey’s multiple comparisons test was used for statistical analysis). **(B)** Comparison of Kavg of different immune cell networks (created using different thresholds (thr)) using network analysis (n = 134 islets from 11 mice) (RM one way ANOVA with Geisser-Greenhouse correction and Tukey’s multiple comparisons test was used for statistical analysis). **(C)** Relationship between the node degrees of different β – immune cell networks obtained using network analysis (different thresholds) (n = 134 islets from 11 mice) (Simple linear regression was used for statistical analysis).

**Supplementary Figure S4. (A)** T-cell density of the islets sorted using T/β cell ratios (insulitis psuedotime) (n = 134 islets from 11 mice) (Non-linear regression curve fit was used for statistical analysis). **(B)** T-cell density curve in A fit using a log scale (n = 134 islets from 11 mice). **(C)** Comparison of insulitis degrees of islets classified as early- (n = 44 islets), intermediate- (n = 46 islets) and late- (n = 44 islets) stage insulitis. **(D)** Insulitis psuedotime in Figure 2B fit using a log scale (n = 134 islets from 11 mice). **(E)** Relationship between the number of β-cells and the number of T-cells in each islet (n = 134 islets). **(F)** Progression of insulitis of different islets in a single mouse sorted using T/β cell ratios (psuedotime) (E-early-stage insulitis, I-intermediate stage insulitis, L – late-stage insulitis). Simple linear regression was used for statistical analysis in B, D and E.

**Supplementary Figure S5. (A)** Schematic representation of the algorithm used to divide an islet into two equal halves. **(B)** Relationship between the proportion of α-cells and the proportion of T-cells in half islets at a distance of at a distance of 20 μm from the islet rim (n = 134 islets) (horizontal division). **(C)** Relationship between the proportion of β-cells and the proportion of T-cells in half islets at a distance of at a distance of 20 μm from the islet rim (n = 134 islets) (vertical division). **(D)** Relationship between the proportion of β-cells and the proportion of T-cells in half islets at a distance of at a distance of 20 μm from the islet rim (n = 134 islets) (horizontal division). Simple linear regression was used for statistical analysis in B, C and D.

**Supplementary Figure S6.** α-cell and T-cell positions of a representative early-stage islet, and 10 seeds of the representative islet randomized using the proposed randomization algorithm

**Supplementary Figure S7. (A)** Experimental and random α-cell – T-cell networks of a representative early-stage islet (positions of α-cells were randomized). **(B)** Comparison of Kavg of experimental and random α-cell – T-cell networks (nwks) in early (n = 44 islets), intermediate (interm) (n = 46 islets), and late (n = 44 islets) stages using network analysis. **(C)** Experimental and random α-cell – macrophage (MLJ) networks of a representative early-stage islet (positions of α-cells were randomized). **(D)** Comparison of Kavg of experimental and random α-cell – macrophage networks in early (n = 44 islets), intermediate (n = 46 islets), and late (n = 44 islets) stages using network analysis. **(E)** Experimental and random α-cell – myeloid (mye) cell networks of a representative early-stage islet (positions of α-cells were randomized). **(F)** Comparison of Kavg of experimental and random α-cell – myeloid cell networks in early (n = 44 islets), intermediate (n = 46 islets), and late (n = 44 islets) stages using network analysis. Paired parametric t tests were used for statistical analysis in B, D and F. See Materials and Methods for detailed description on randomization and network analysis.

**Supplementary Figure S8. (A)** Comparison of Kavg of experimental and random α – T cell networks in early- (n = 44 islets), intermediate- (n = 46 islets) and late – (n = 44 islets) stage islets using network analysis (different thresholds (thr)). **(B)** Comparison of Kavg of experimental and random α-cell – macrophage (Mφ) networks in early- (n = 44 islets), intermediate- (n = 46 islets) and late – (n = 44 islets) stage islets using network analysis (different thresholds). **(C)** Comparison of Kavg of experimental and random α – myeloid (mye) cell networks in early- (n = 44 islets), intermediate- (n = 46 islets) and late – (n = 44 islets) stage islets using network analysis (different thresholds). Paired parametric t tests were used for statistical analysis in A-C.

**Supplementary Figure S9.** α-, β- and T- cell positions of 10 randomly picked early-stage islets.

**Supplementary Figure S10. (A)** Progression of insulitis in different islets, sorted using T/β cell ratios (psuedotime) (n = 134 islets from 11 mice). **(B)** Comparison of Kavg of small sized experimental and random α – T cell, and experimental and random α-cell – macrophage (Mφ) networks in early-, intermediate- and late – stage islets using network analysis. **(C)** Comparison of Kavg of medium sized experimental and random α – T cell, and experimental and random α-cell – macrophage (Mφ) networks in early-, intermediate- and late – stage islets using network analysis. **(D)** Comparison of Kavg of large sized experimental and random α – T cell, and experimental and random α-cell – macrophage (Mφ) networks in early-, intermediate- and late – stage islets using network analysis. Paired parametric t tests were used for statistical analysis in B-D.

**Supplementary Figure S11. (A)** Comparison of Kavg of α-linked β-cell – and non-α-linked β-cell – T cell networks in early- (n = 44 islets), intermediate- (n = 46 islets) and late- (n = 44 islets) stage islets using network analysis (different thresholds (thr)). **(B)** Comparison of Kavg of α-linked β-cell – and non-α-linked β-cell – macrophage (Mφ) networks in early- (n = 44 islets), intermediate- (n = 46 islets) and late – (n = 44 islets) stage islets using network analysis (different thresholds). **(C)** Comparison of Kavg of α-linked β-cell – and non-α-linked β-cell – myeloid (mye) cell networks in early- (n = 44 islets), intermediate (n = 46 islets) and late- (n = 44 islets) stage islets using network analysis (different thresholds). Wilcoxon matched-pairs signed rank test was used for statistical analysis in A-C.

**Supplementary Figure S12: (A)** Progression of insulitis in different islets, sorted using T/β cell ratios (psuedotime) (n = 134 islets from 11 mice). **(B)** Comparison of Kavg of small sized α-linked β- cell – and non-α-linked β-cell – T cell, and experimental and random α-linked β-cell – and non-α- linked β-cell – macrophage (Mφ) networks in early-, intermediate- and late – stage islets using network analysis. **(C)** Comparison of Kavg of medium sized α-linked β-cell – and non-α-linked β- cell – T cell, and α-linked β-cell – and non-α-linked β-cell – macrophage (Mφ) networks in early-, intermediate- and late – stage islets using network analysis. **(D)** Comparison of Kavg of large sized α-linked β-cell – and non-α-linked β-cell – T cell, and α-linked β-cell – and non-α-linked β-cell – macrophage (Mφ) networks in early-, intermediate- and late – stage islets using network analysis. Wilcoxon matched-pairs signed rank test was used for statistical analysis in B-D.

