## Supplemental Figure S1 for "Network approach reveals preferential T-cell and macrophage association with α-linked β-cells in early stage of insulitis in NOD mice"

| Cell types | M1 | M2 |
|---|---|---|
| **CD3 and CD11c** | 0.20347907 ± 0.002117053 | 0.426356977 ± 0.003970354 |
| **CD3 and F4/80** | 0.270915116 ± 0.002479906 | 0.253376744 ± 0.003874686 |
| **CD11c and F4/80** | 0.513327907 ± 0.002275661 | 0.208355814 ± 0.001571986 |
