## Supplemental Figure S2 for "Network approach reveals preferential T-cell and macrophage association with α-linked β-cells in early stage of insulitis in NOD mice"

| Mouse no | Age (weeks) | Highest blood glucose recording (mg/dL) | No of intact islets analyzed in the cross section | Range of insulitis degrees |
|---|---|---|---|---|
| **1** | 19.1 | 154 | 2 | 1.371 – 4.029 |
| **2** | 22.3 | 220 | 11 | 0.257 – 7.800 |
| **3** | 20.4 | 168 | 5 | 0.721 – 3.583 |
| **4** | 16.1 | 154 | 6 | 0.510 – 4.091 |
| **5** | 19.4 | 179 | 14 | 0.065 – 5.692 |
| **6** | 20.1 | 190 | 13 | 0.371 – 15.143 |
| **7** | 16.7 | 151 | 28 | 0.0217 – 27.500 |
| **8** | 16.7 | 195 | 19 | 0.136 – 4.817 |
| **9** | 17 | 135 | 11 | 0.185 – 7.610 |
| **10** | 21.7 | 180 | 3 | 0.381 – 3.000 |
| **11** | 21.7 | 157 | 22 | 0.0595 – 3.387 |
