## Supplementary figures and images for "Network approach reveals preferential T-cell and macrophage association with α-linked β-cells in early stage of insulitis in NOD mice"

### Supplemental Figure S3

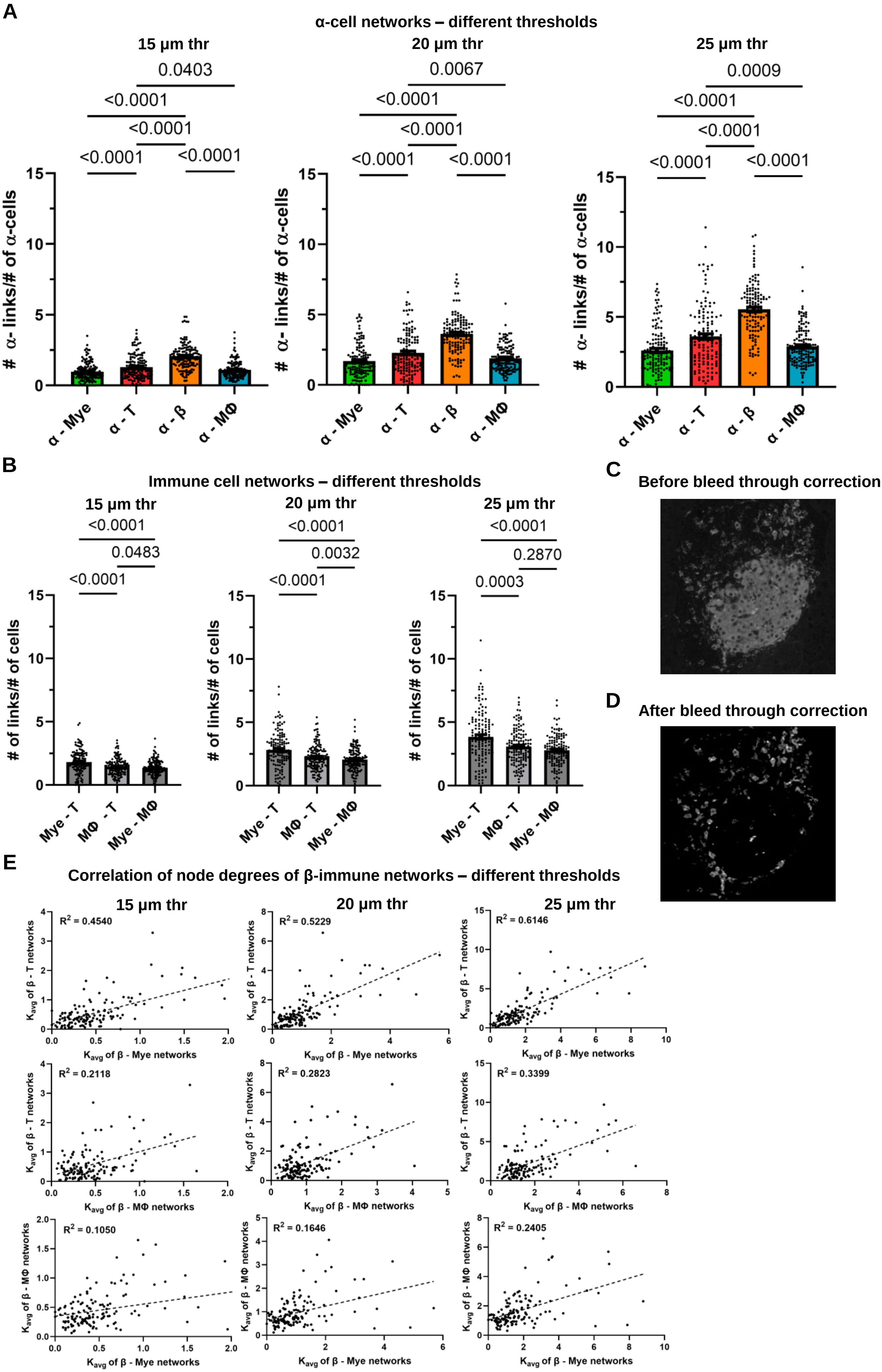

### Supplemental Figure S4

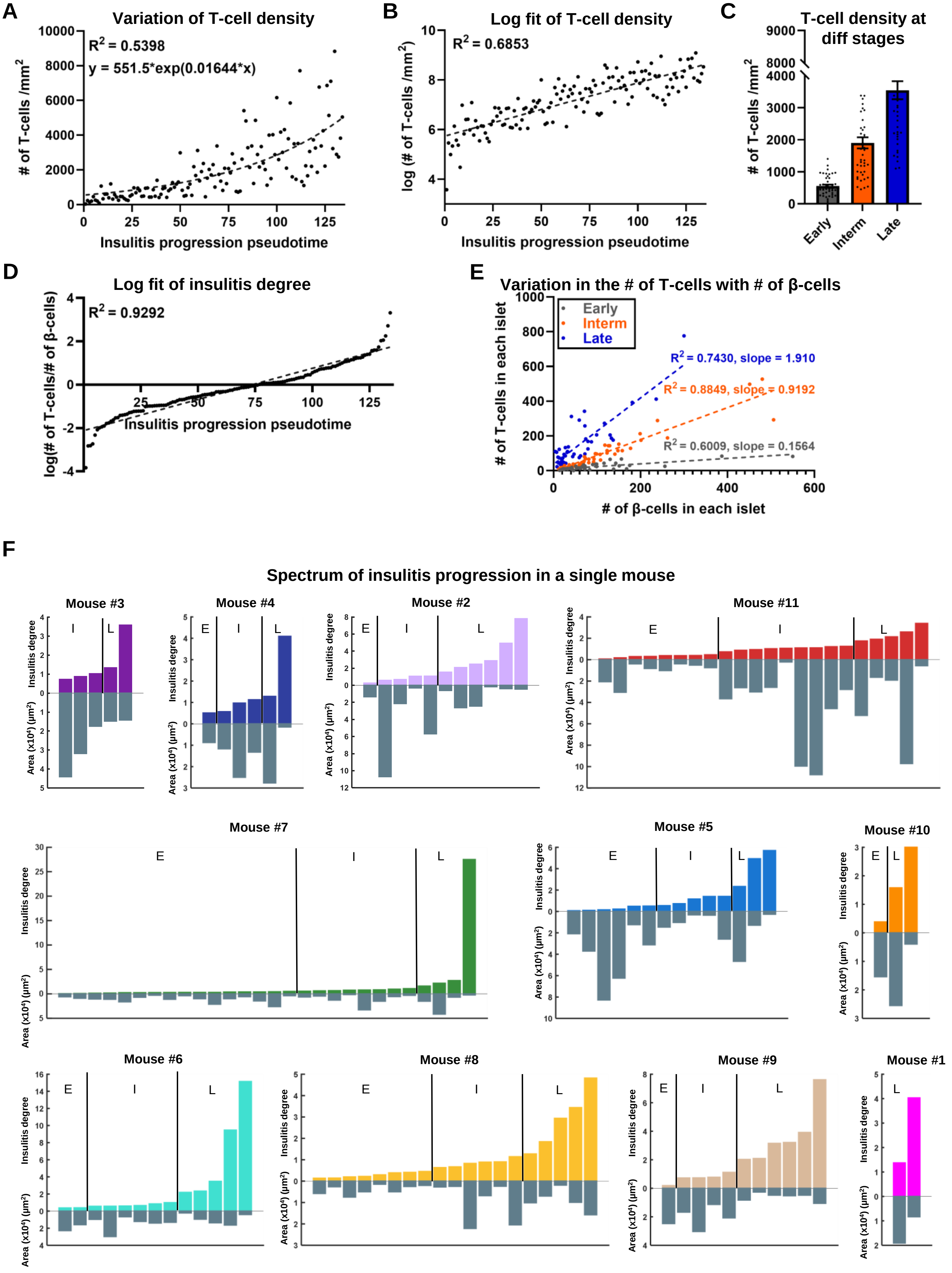

### Supplemental Figure S5

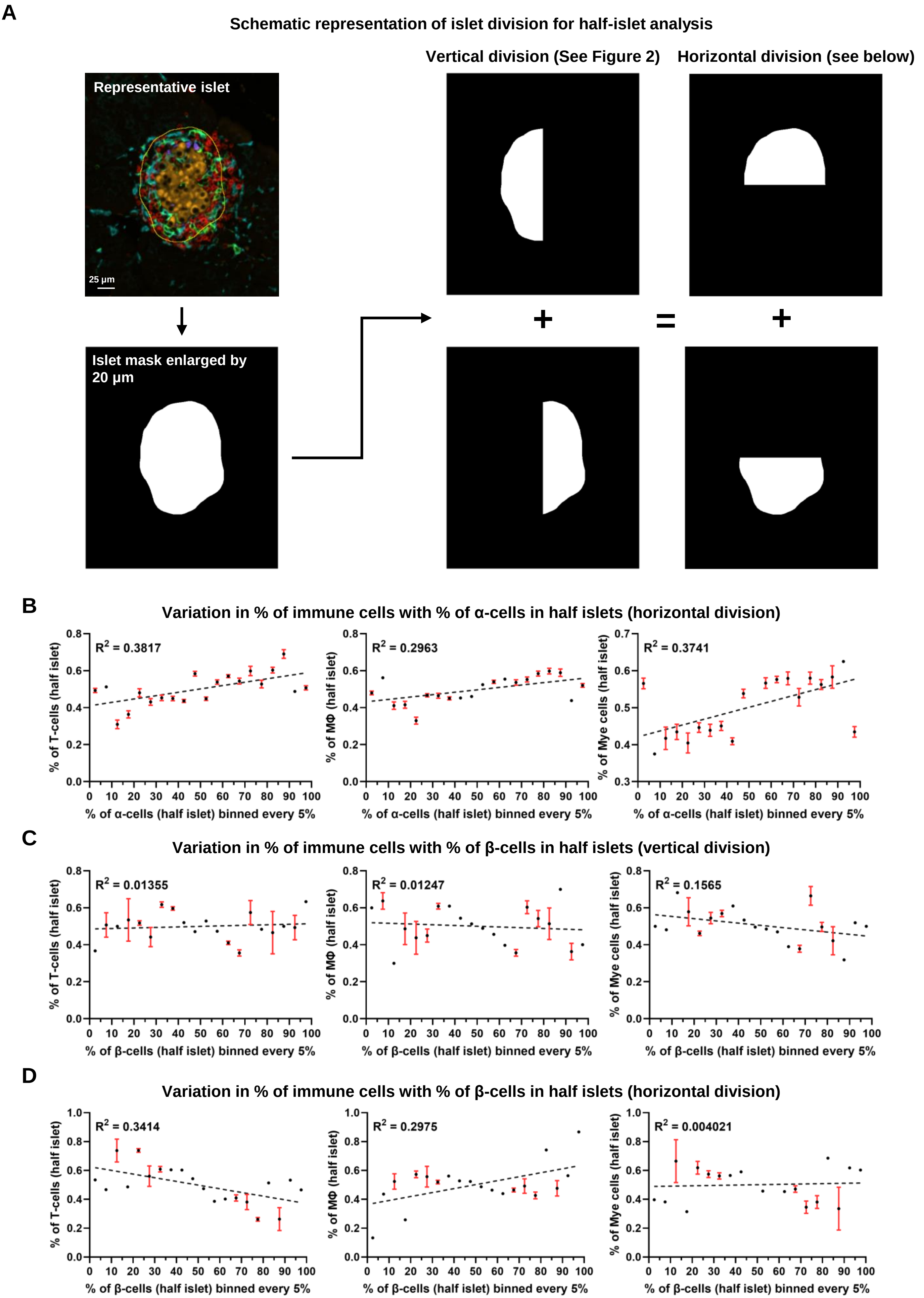

### Supplemental Figure S6

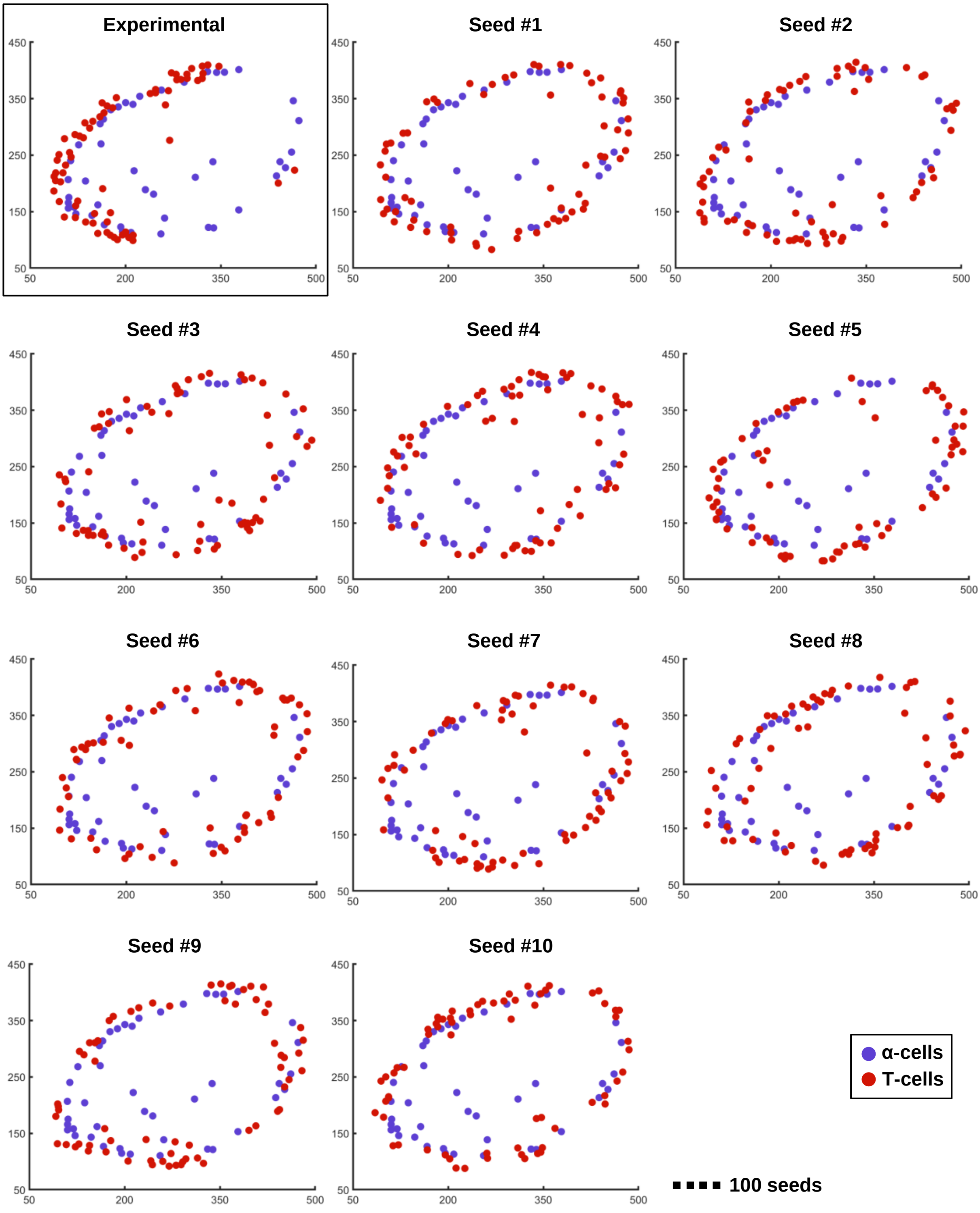

### Supplemental Figure S7

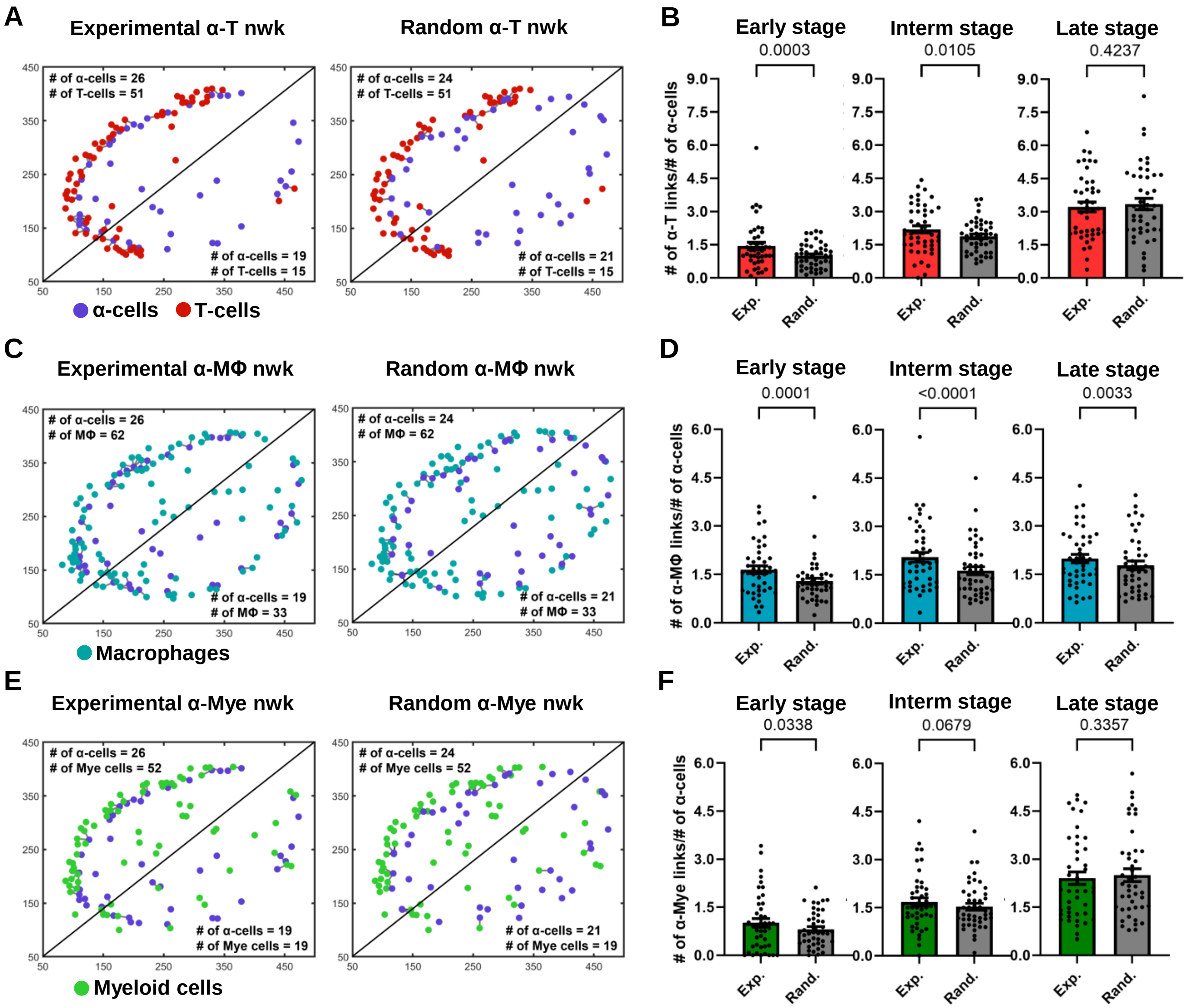

### Supplemental Figure S8

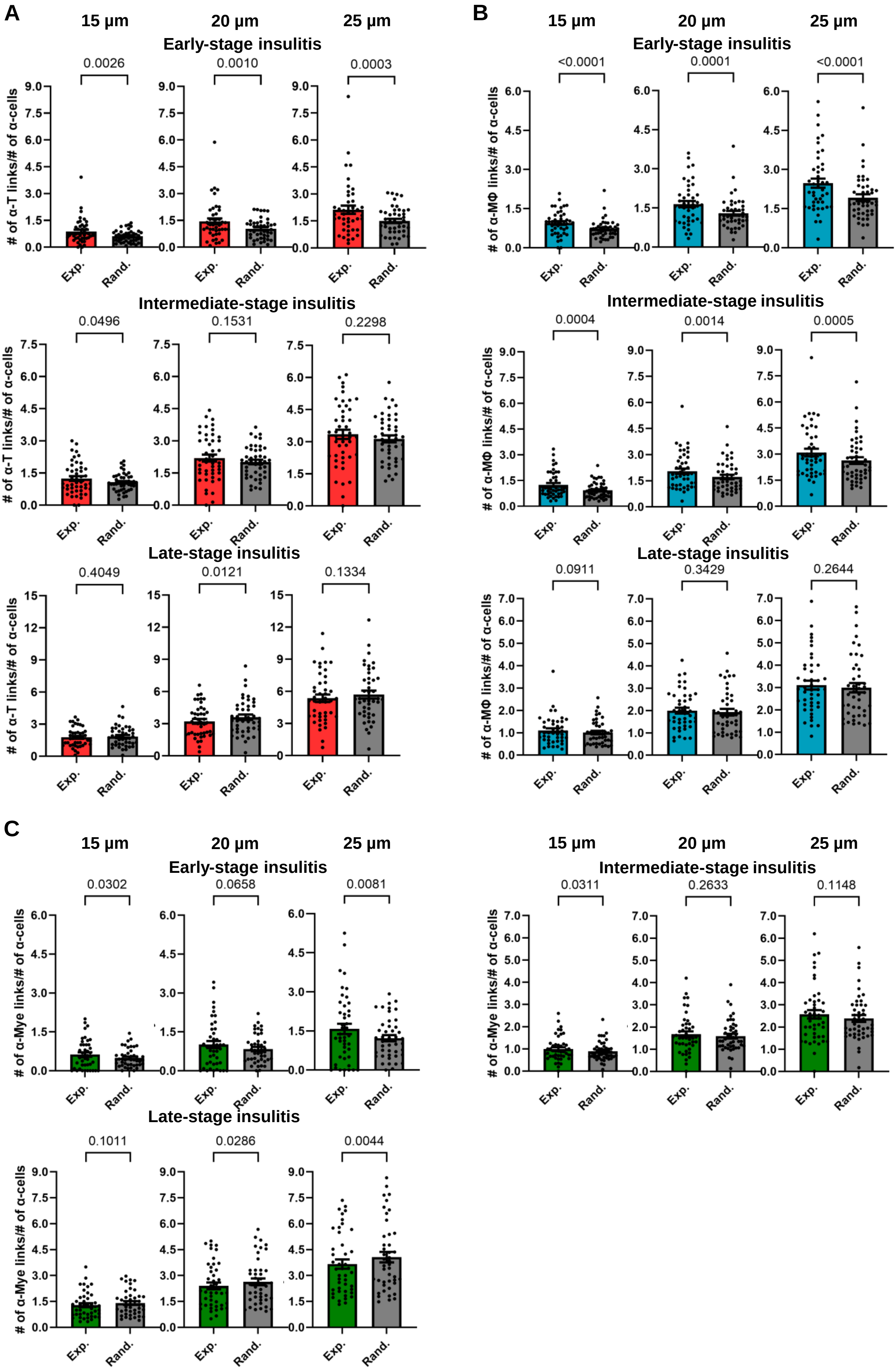

### Supplemental Figure S9

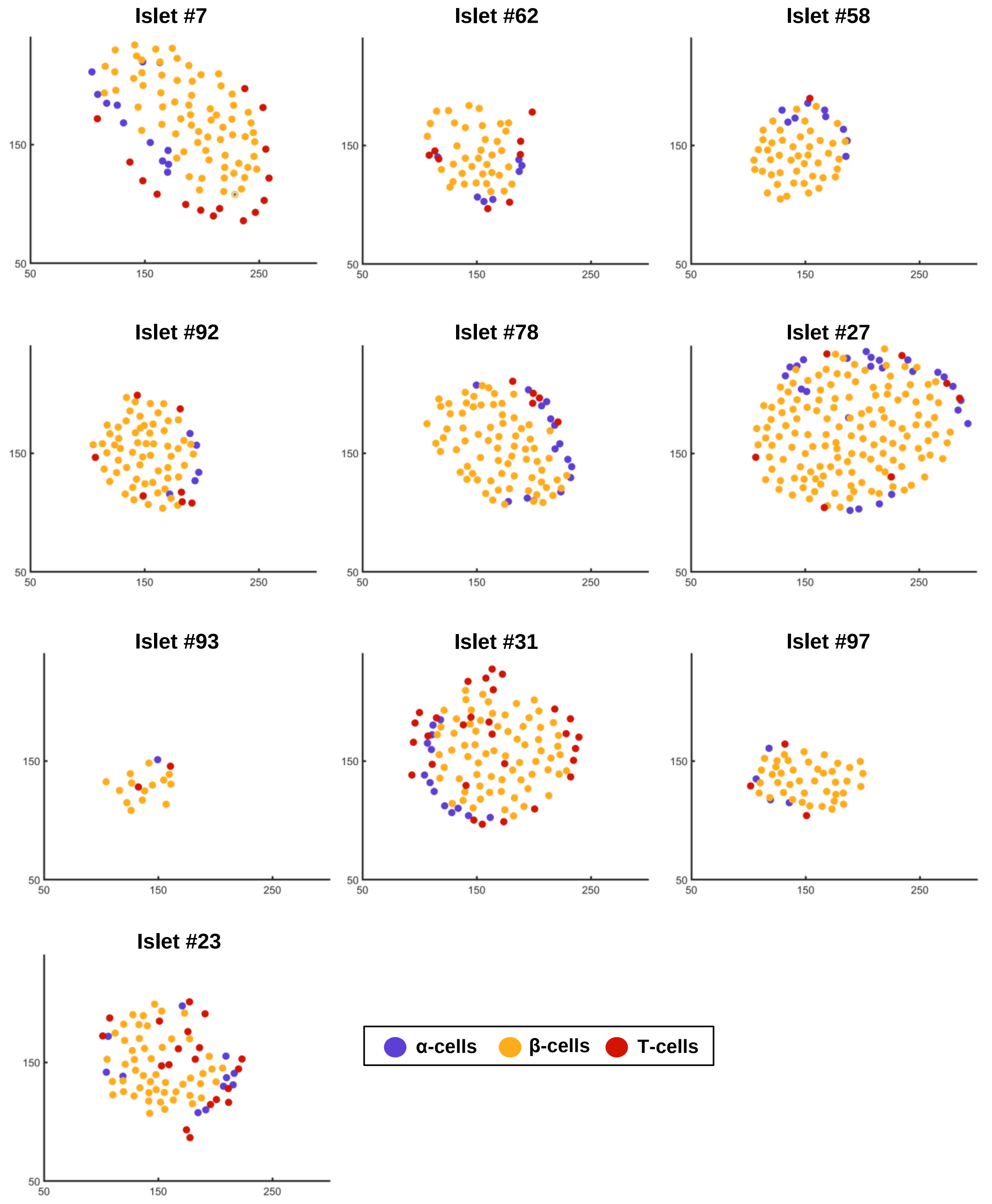

### Supplemental Figure S10

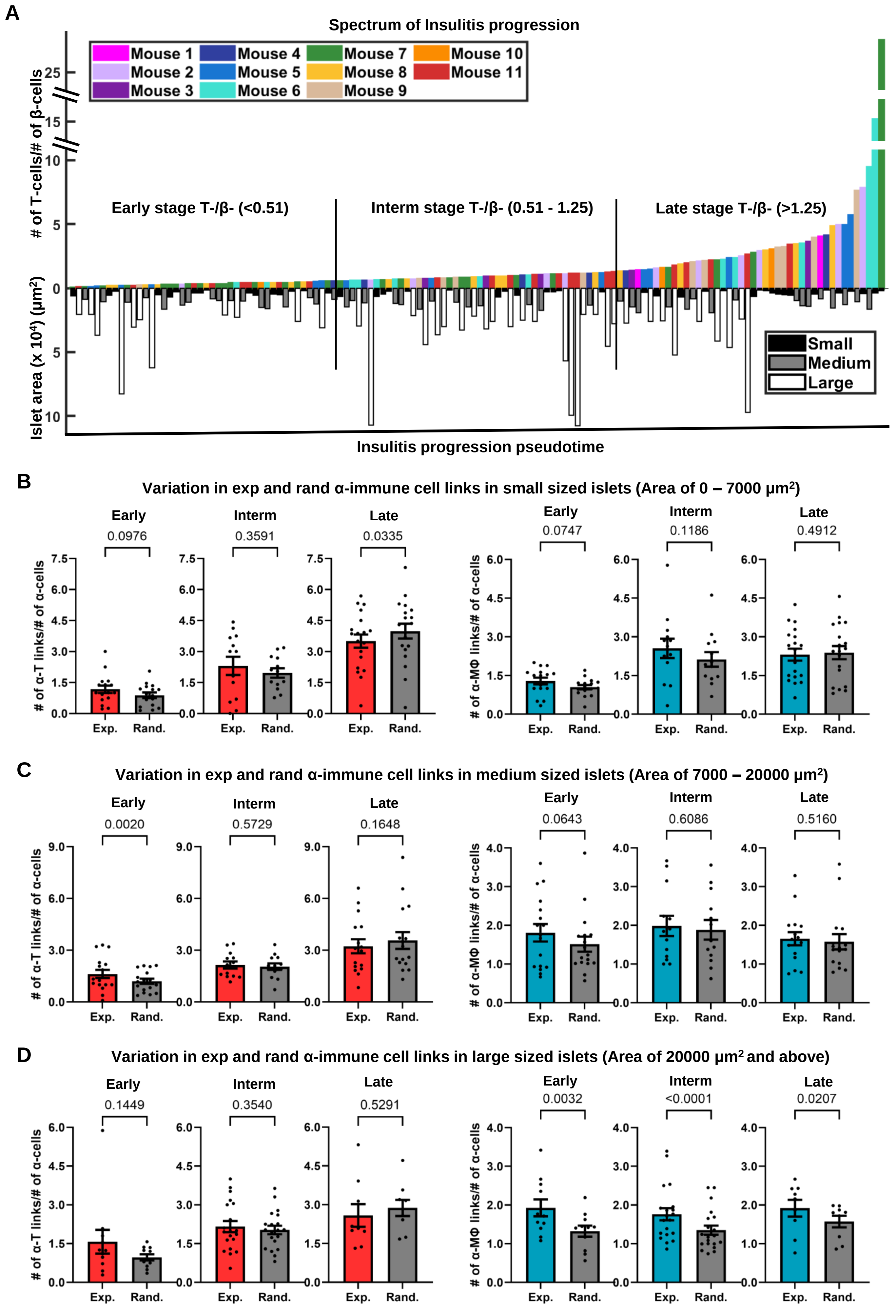

### Supplemental Figure S11

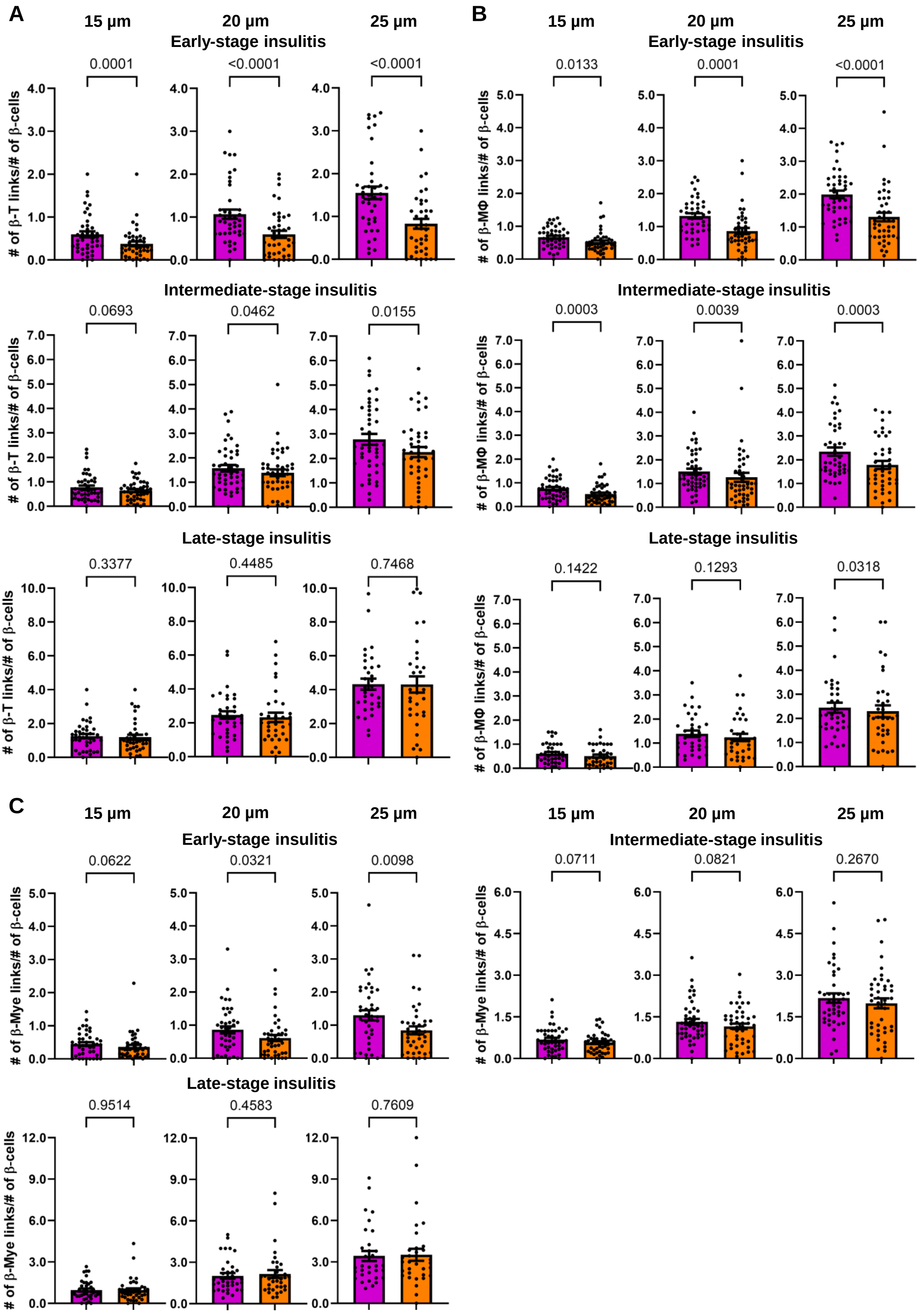

### Supplemental Figure S12

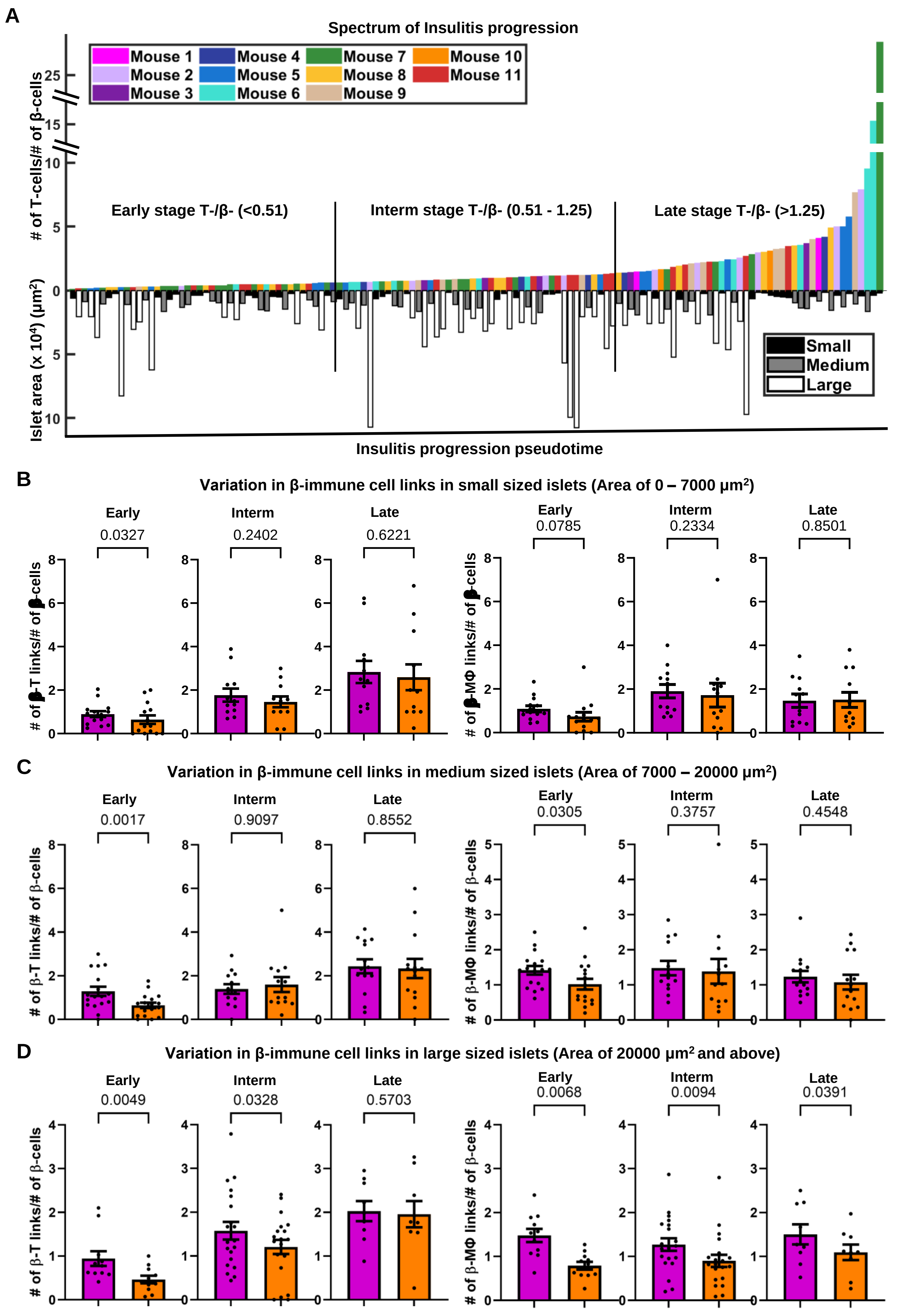
